# Achievement-Based Differences in Cognitive-Emotional Interplay During Classroom Learning: A Multimodal Analysis with Experience Sampling Method

**DOI:** 10.1101/2025.01.23.634609

**Authors:** Xiaobo Liu, Lan Fu, Zihao Yang, Zheng Zhang, Yu Zhang

## Abstract

**Background:** The relationship between cognition and emotion during learning is crucial for educational theories and has been under debate for a long time, from the ancient philosophy to current scientific analyses. However, the relationship may not be uniform and may vary across diverse individual groups.

**Aims:** Therefore, this study aims to explore whether the relationship between cognition and emotion varies across different student groups. Furthermore, it examines whether neurophysiological evidence can provide insights into such group differences.

**Sample:** The study involved 22 high school students (11 high-performing and 11 low-performing) over 13 math lessons. Surveys were administered three times during each lesson, yielding a multilevel dataset with 777 samples.

**Methods:** The experience sampling method was employed to gather data on cognitive and emotional states periodically, as well as their regulations. Simultaneously, multimodal neurophysiological data, including electroencephalograph (EEG), electrodermal activity (EDA), and photoplethysmography (PPG), were collected during classroom learning to analyze the relationship further.

**Results:** Cognitive state, cognitive regulation, emotional state, and emotional regulation were highly correlated overall. In high-performing students, the association was higher than that in low-performing students, especially at the level of within-subject repeated-measures. Multimodal data revealed the common neurophysiological basis of the correlated variation between cognition and emotion, and highlighted different neurophysiological patterns between high and low achievers.

**Conclusions:** The joint evidence from the experience sampling survey and neurophysiological measures reveals that the cognitive-emotional relationship differs by academic achievement levels, with high achievers revealing a more consistent variation between cognition and emotion. Such co-variation shares a common neurophysiological pattern with features associated with high student-teacher interaction and high concentration, further explaining the group differences.

## 1. Introduction

The relationship between cognition and emotion has been long debated since the early days of western philosophy (Lyons, 1999; Pessoa, 2008). One perspective argues that cognition and emotion are distinct and separate, while the other asserts that they are inseparable and mutually dependent.

This fundamental debate has shaped the core assumptions of educational theories. The debate spans multiple disciplines including philosophy, psychology, and neuroscience. The philosophical debate on the relationship between cognition and emotion centers on separation versus integration. For instance, Plato regarded emotion as the lower part of the rational soul and emphasized the need to control it, while Aristotle saw emotion as a natural part of human life that could be guided by reason to support virtuous actions (Lazarus, 1999). The debate in psychology centers on understanding the mechanisms of cognition and emotion. Cognitive appraisal theory proposes that emotion originates from cognitive processes (Smith & Ellsworth, 1985), while affective primacy theory emphasizes the independence of emotion (Zajonc, 1984). Neuroscientific discussions focus on the physiological structures corresponding to the functions of cognition and emotion. Early functional localization theories assumed that cognition and emotion function at distinct brain regions. However, recent neuroimaging evidence challenges this view, indicating that many brain regions are jointly involved in emotional and cognitive tasks (Pessoa, 2008). Despite the longstanding debate, there is still no conclusive consensus on this issue.

While these studies attempt to analyze the general relationship between cognition and emotion, there are very few discussions on the group differences in this relationship. In the classroom, cognitive and emotional experiences can be shaped by a range of group differences, including learning strategies and demographic factors. Extensive research has already shown substantial group differences among learners in cognitive strategies or emotional responses, through various research approaches (Hendricks & Buchanan, 2016; Price, 2004). However, most empirical studies are conducted in laboratory settings, whereas the classroom learning is diverse and dynamic. In classroom learning, the variance and covariance of cognition and emotion at the within-subject level may reflect how individuals engage with cognition and emotion across different learning activities. Examining the cognitive-emotional relationship at the within-subject level offers deeper insights into its dynamics over time and the potential underlying group differences.

Therefore, this study aims to explore the group differences in the relationship between cognition and emotion. Specifically, academic performance is adopted to categorize students into two groups. Two research questions are proposed. **RQ1:** *Do high-performing and low-performing students differ in the relationship between cognition and emotion?* To address RQ1, this study employed a repeated-measures design using the experience sampling method, analyzing the cognitive-emotional relationship at both between-subject and within-subject levels. **RQ2:** *Do two groups of students differ in the shared neurophysiological representations of cognition and emotion?* In neuroscience studies, when cognition and emotion share neurophysiological representations, this indicates that these two processes are interrelated at the neural level. Therefore, analyzing the shared representations can deepen our understanding of the cognitive-emotional relationship at the neural level, supporting findings from subjective measures.

## 2. Literature Review

### 2.1 Psychological discussions on the cognitive-emotional relationship

Recent psychological discussions argued that there is a complex interaction between basic cognitive and emotional functions (Liu et al., 2009). Emotion influences working memory performance through a motivation-based approach, as it mobilizes cognitive resources by driving motivation (Yüvrük et al., 2020). In a reciprocal manner, cognition also plays an important role in emotional processes. When individuals focus their attention on the positive or negative aspects of a stimulus, this allocation of attention directly influences their self-report emotional state (Freund & Keil, 2013). Similarly, the exposure effect suggests that by adjusting the frequency of exposure to a stimulus, participants’ preferences can be influenced (Montoya et al., 2017). The evidence suggests a complex interaction between cognition and emotion at the level of basic functions.

Moreover, both cognition and emotion are dynamically regulated, with the regulatory process involved in their interaction. Cognitive reappraisal, as a core strategy for emotional regulation (Brockman et al., 2017), is inherently a cognitive process. This strategy involves altering the generation or intensity of emotions by re-evaluating the meaning of a situation, effectively transforming the internal emotional experience. Cognitive control has also been shown to effectively reduce negative affect, as evidenced by the activation of associated brain regions (Ochsner et al., 2002). Emotional regulation not only involves cognitive processes but also impacts cognition in return. A study analyzed the effects of two emotional regulation strategies, expressive suppression and cognitive reappraisal, on memory (Gross, 2002). Expressive suppression negatively impacts memory, whereas the reappraisal strategy does not. These findings suggest that the regulatory process plays an important role in the relationship between cognition and emotion. To gain a deeper understanding of this relationship, cognitive and emotional regulation should be considered in addition to cognitive and emotional states.

In educational psychology, theoretical studies focus on how cognition and emotion jointly support learning. For example, the investment of cognition and emotion in learning is conceptualized as cognitive and emotional engagement (Fredricks et al., 2004). These two aspects, along with behavioral engagement, interact with and influence each other, collectively promoting student learning (Pietarinen et al., 2014). Another influential theory, the control-value theory, focuses on how emotions arise and influence learning (Pekrun, 2006). This theory posits that achievement emotions arise from appraisals of control and value, which are closely linked to cognitive processes. When students positively appraise their sense of control and the value of a task, they are more likely to experience positive emotions such as joy and hope. These emotions, in turn, enhance motivation and ultimately improve learning outcomes (Wu & Yu, 2022). These theories, from a meso-level perspective, establish a connection between the cognitive-emotional relationship and effective learning.

Empirical studies suggest that self-reports of cognition and emotion exhibit strong associations, supporting their interconnectedness. In a three-year dataset examining the cognitive and emotional engagement of high school students, Li and Lerner (2013) found that cognitive and emotional engagement not only showed high cross-sectional correlations (on average r = .55) but also demonstrated cross-lagged effects in the second wave. Similarly, Obergriesser and Stoeger (2020) identified a unidirectional effect of enjoyment on the effective use of learning strategies. An analysis of MOOC discussions also revealed strong associations between cognitive and emotional engagement (on average r = .48), with the highest correlation found between positive emotions and constructive cognition (Liu et al., 2022).

Although the cognitive-emotional relationship in the learning process has been widely discussed, there is little research addressing the potential group differences in this relationship. Cognitive and emotional experiences in learning are abundant and various (He et al., 2024), suggesting variety in the relationship between cognition and emotion. Such variety may be influenced by previous experience, cultural factors, or academic achievement. For example, the relationship between achievement and emotions is found to be mediated by self-concept, with high-performing students exhibiting stronger self-concepts, more enjoyment, and less math anxiety (Van der Beek et al., 2017). In a longitudinal study, reciprocal effects were found between emotions and academic performance, highlighting the impact of achievement on academic emotions (Pekrun et al., 2017). These studies suggest that the cognitive-emotional relationship may vary depending on multiple group factors.

Furthermore, when analyzing potential group differences, a dynamic perspective may offer a valuable viewpoint. Research has shown that students’ cognition and emotions fluctuate over the course of a semester or even within a few weeks (Faloughi & Herman, 2021; Xie, Heddy, & Greene, 2019), highlighting the temporal variability of learning experiences. The within-subject temporal dynamics of engagement were further found to be associated with perceived challenge, skill level, and emotional states during learning activities (Schneider et al., 2016). Therefore, analyzing the temporal dynamics within individuals may provide valuable insights into how the cognitive-emotional relationship varies across different student groups.

### 2.2 Neural evidence on the cognitive-emotional relationship

To support cognitive and emotional activities, both the central nervous system and the peripheral nervous system are involved. Neuroscientific research has extensively explored neural activities in cognitive and emotional processes. These studies, on the one hand, help us understand the mechanisms underlying the relationship between cognition and emotion. On the other hand, these studies provide measurement tools, enabling us to observe the dynamic changes in cognition and emotion in more complex contexts.

Human cognitive architecture encompasses various functions, including attention, working memory, and long-term memory, which are associated with brain activity in regions such as the prefrontal cortex and hippocampus. The prefrontal cortex serves as the core area for executive functions particularly for working memory and inhibitory control (Moriguchi & Hiraki, 2013). The hippocampus is responsible for encoding episodic and spatial memory, participating in the consolidation of short-term memory into long-term memory (Bird & Burgess, 2008; Schapiro et al., 2019). Emotional processing involves areas of the hypothalamus and amygdala. As a key region connecting multiple levels of the nervous system, the hypothalamus is considered the center for emotional responses (Rolls, 1990). The amygdala is involved in the perception, evaluation, and regulation of emotions, particularly the experience of fear (Gallagher & Chiba, 1996).

In research on the neural mechanisms of cognition and emotion, emotional activities are often found to engage brain regions typically associated with cognitive processes, and vice versa. For example, the hippocampus was initially considered part of “the emotional brain”, but it is now no longer considered critically linked to affect (Pessoa, 2008). The activation of the hippocampus in response to emotional stimuli can now be explained by the amygdala-hippocampal interaction. This suggests that the amygdala modulates the encoding and storage of hippocampal-dependent memories, while the hippocampus influences the amygdala response when encountering emotional stimuli (Phelps, 2004; Richardson et al., 2004). The orbitofrontal cortex, a subregion of the prefrontal cortex, is also involved in the interaction between cognition and emotion. This area is thought to process reward information, playing a key role in guiding decisions (Wallis, 2007). A study on decision-making indicates that the orbitofrontal cortex is activated both when inhibiting emotional interference and when integrating emotional information into decision-making (Beer et al., 2006). Based on the evidence that some brain regions are both “cognitive” and “affective”, Pessoa (2008) proposed that cognition and emotion are not processed separately in the brain but are dynamically integrated through interconnected networks.

In addition to evidence from the central nervous system, findings from the peripheral nervous system also shed light on the cognitive-emotional relationship. A series of physiological responses, such as accelerated heartbeat, rapid breathing, and sweating, were initially attributed to emotions or stress. The mechanism behind this phenomenon is that when students encounter emotional or stress-related stimuli, the hypothalamus sends signals to regulate the peripheral nervous system, particularly the sympathetic nervous system, preparing the body for an appropriate response (Sequeira et al., 2009). Recent studies have found that cognitive activities can also trigger peripheral physiological responses (Critchley et al., 2013), such as cognitive load triggering electrodermal activity (Setz et al., 2009). This indicates that peripheral responses are linked to both cognitive and emotional activities, highlighting their potential for uncovering the relationship between cognition and emotion.

Portable biosensors offer new opportunities for analyzing neural activities during complex real-world tasks, including classroom learning. Neurophysiological signals that can be collected using wearable devices include electroencephalograph (EEG), photoplethysmography (PPG), and electrodermal activity (EDA). The band powers of the EEG signal have been successfully applied to predict both attention (Liu et al., 2013) and emotional states (Kostyunina & Kulikov, 1996). Among various spectral bands, alpha reflects the brain’s “wakefulness” state (Klimesch et al., 2007), and theta is associated with cognitive control (Cavanagh & Frank, 2014). PPG signals can be used to evaluate heart rate and pulse rate variability, which reflect the cardiovascular system’s response to external stimuli. These features have been found to be related to emotional arousal (Udovičić et al., 2017) or cognitive load (Qi et al., 2020). Analysis of EDA signals typically decomposes them into two components: skin conductance response (SCR) and skin conductance level (SCL), which reflect rapid changes and slower trends in skin conductance, respectively (Boucsein, 2012). The two components are associated with both cognitive and emotional states (Paletta et al., 2015). In addition to individual metrics, researchers have also found that the inter-subject correlation (ISC) is a promising indicator, reflecting the degree to which different individuals exhibit similar neural activities when exposed to the same type of stimuli. Therefore, ISC can somehow reflect learners’ cognitive and emotional levels without labeling the complex stimuli in learning. For example, studies have shown that ISC is associated with academic performance (Dikker et al., 2017), as well as students’ cognitive and emotional engagement (Gao et al., 2020).

## 3. Methods

### 3.1 Research design

This study was conducted at a high school in Beijing and involved 22 Grade-10 students (9 girls and 13 boys, aged 15-16 years) across 13 math lessons, as illustrated in Figure 1a. For each 40-minute lesson, the ESM survey was administered three times, dividing the lesson into sessions of 10-15 minutes each. To standardize the time duration, students were asked to rate their feelings based on their experiences over the past 10 minutes. During the classroom learning, multimodal neurophysiological data were simultaneously collected. A total of 777 samples formed the basis for the analysis. The repeated-measures dataset allows for analysis at both the within-subject and between-subject levels.

**Figure 1.**
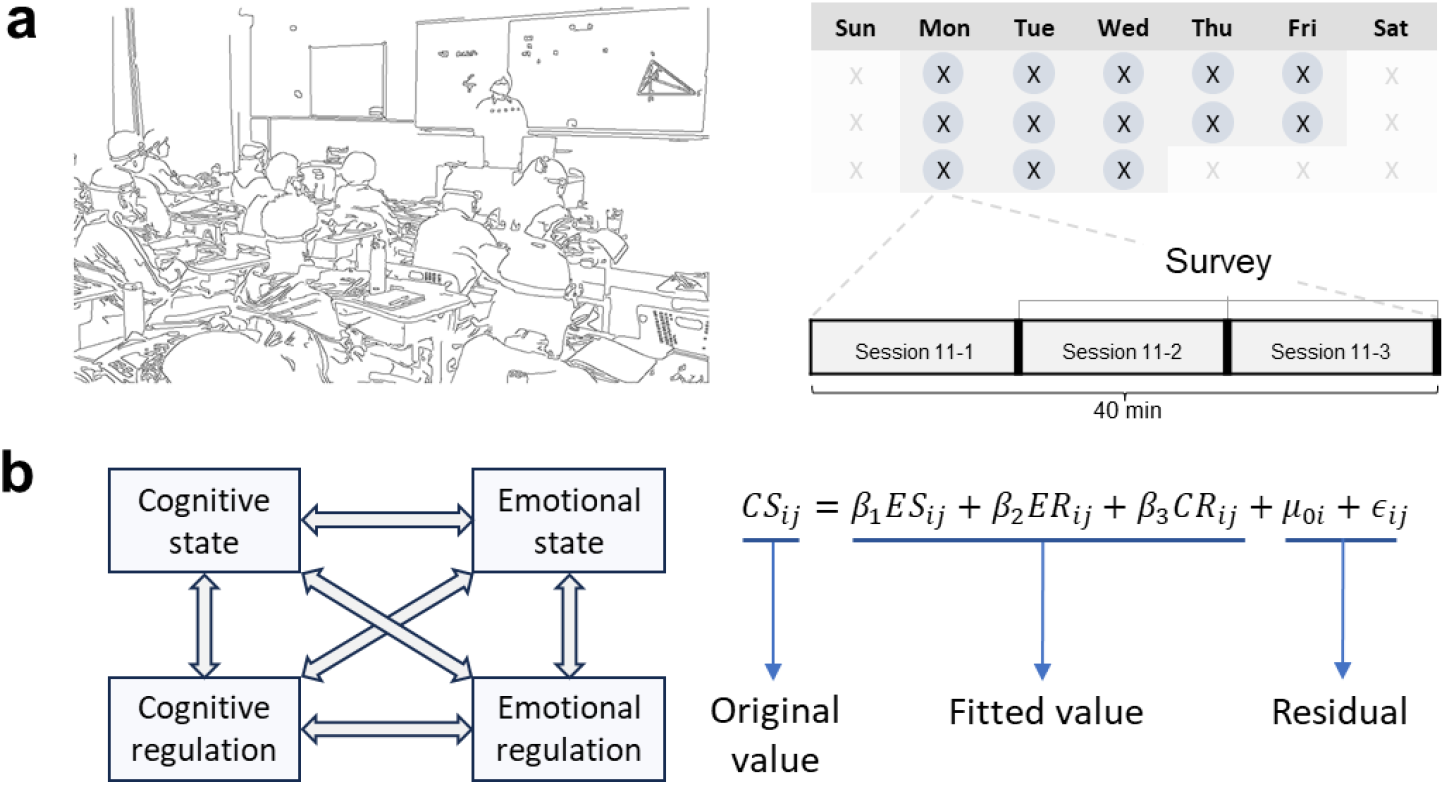
Research framework. a. Experimental design. Data were collected in a real-world learning environment, involving 22 students across 13 lessons. b. Analytical framework. The Pearson correlation between the variables was first examined, and a multilevel regression model was then constructed to decompose the interrelated variables into fitted values and residuals, followed by a neurophysiological analysis.

To examine how the indices of emotion and cognition are influenced by group differences, the 22 students in this study were divided into high-performing and low-performing groups based on their scores in a pre-experiment exam. The top 11 students were classified as high-performing, while the remaining students were considered low-performing. Considering the data structure of repeated measurements, the following random effect model was used for analyzing the differences in survey data between different groups:

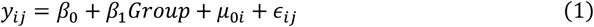

where *y*_*ij*_ represents the survey data for student i in session j. The coefficient *β*_1_ represents the group difference, and *β*_0_, *μ*_0*i*_, and ϵ_*ij*_ are the intercept, the between-person residual, and the within-person residual, respectively.

This study then investigated the relationship between cognitive state (CS, specifically attention), cognitive regulation (CR), emotional state (ES, specifically positive emotion), and emotional regulation (ER) (see Figure 1b). For simplicity, we first analyzed the Pearson correlations among these four variables. Next, we constructed a multilevel regression model to analyze how each of the four variables is explained by the other three variables. Specifically, equation (2) shows the model where emotional state is explained by the other three variables:

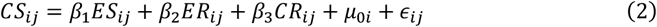

where *CS*_*ij*_, *CR*_*ij*_, *ES*_*ij*_, and *ER*_*ij*_ are the variables of cognitive state, cognitive regulation, emotional state, and emotional regulation, respectively. In this model, the within-group, between-group, and overall R^2^ values will be reported, illustrating the cognitive-emotional relationship at different levels.

Following equation (2), the value of cognitive state can be decomposed into fitted values and residuals, representing explained and unexplained contributions, respectively. Operationally, the fitted values were computed through equation (3), and the residuals were the difference between the original values and the fitted values. Taking cognitive state as an example, its fitted value is:

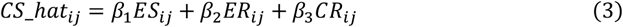

where *CS_hat*_*ij*_ represents the fitted value. In this way, we represent the relationship between cognition and emotion using two entities: fitted values and residuals. Next, a multimodal neurophysiological analysis was conducted to explore the differences in neural bases between individual groups and between fitted values and residuals, as illustrated in equation (4):

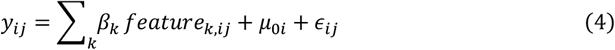

where *y*_*ij*_ represents one of the original values, fitted values, or residuals, and *feature*_*k,ij*_ represents one of the multimodal neurophysiological features, which will be described in detail in the section 3.3 *Neurophysiological data collection and processing*.

### 3.2 Experience Sampling Survey

The ESM survey was designed to capture cognitive and emotional states, as well as their regulation. The scale items used in this study are detailed in Table 1. Specifically, the measurement of cognitive and emotional states primarily involves the concepts of attention and positive emotion. The items “*I was focused*” (Ben-Eliyahu et al., 2018; Lam et al., 2014; Reeve & Tseng, 2011) and “*My emotions were positive*” (Barrett & Russell, 1998) were included, as they are commonly used in ESM studies (Xie, Heddy, & Vongkulluksn, 2019). Items related to self-regulation were also included: “*I had clear learning goals and flexibly adapted my learning strategies to achieve them*” for cognitive regulation (Greene, 2015) and “*I always sensed and actively adjusted my emotions*” for emotional regulation (Boemo et al., 2022). All items were reported with a 5-point Likert scale, ranging from “strongly disagree” to “strongly agree”.

**Table 1.** Scale items.

| Type | Items |
| --- | --- |
| Cognitive state | *I was focused.* |
| Cognitive regulation | *I had clear learning goals and flexibly adapted my learning strategies to achieve them.* |
| Emotional state | *My emotions were positive.* |
| Emotional regulation | *I always sensed and actively adjusted my emotions.* |

To validate the effectiveness of the experience sampling method survey (ESM), we conducted an additional survey (**recall method survey**) using a full questionnaire containing the four items, administered to a larger sample prior to the experiment. In the recall method survey, students reflected on their previous class session and reported their scores for cognitive and emotional dimensions. The full scales of cognitive state, and cognitive regulation, emotional state, emotional regulation each passed reliability and validity testing, confirming their effectiveness. Therefore, items from these scales were selected for the experience sampling method survey.

### 3.3 Neurophysiological data collection and processing

Portable biosensors were employed to collect students’ neurophysiological signals, including a head belt (Brainno, SOSO H&C, South Korea) collecting EEG signals at the Fp1 and Fp2 channels at 250 Hz, and a wristwatch (Psychorus, China) collecting EDA and PPG signals at the left wrist, with the sampling rate of 40 Hz and 20 Hz, respectively. These devices have been demonstrated to be effective in real-classroom settings with minimal disruption to learning (Chen et al., 2023; Zhang et al., 2018). The neurophysiological data collection was approved by the Institutional Review Board (IRB) of the Department of Psychology, Tsinghua University. Prior to the experiment, all participants and their legal guardians signed consent forms, and the students received training on how to properly wear these devices.

The EEG raw signals were sequentially processed using robust detrending (de Cheveigné & Arzounian, 2018), band-pass filtering, and ocular artifact removal (Kanoga et al., 2019) to eliminate potential artifacts (Chen et al., 2023). To mitigate the effects of unrecognized artifacts, 5-second epochs with peaks exceeding ±150 μV were excluded. For the remaining epochs, absolute band powers of ***delta*** (1 Hz - 4 Hz), ***theta*** (4 Hz - 8 Hz), ***alpha*** (8 Hz - 13 Hz), ***beta*** (13 Hz - 30 Hz), and ***gamma*** (30 Hz - 45 Hz) were computed. Additionally, the relative band powers (***delta_rel, theta_rel, alpha_rel, beta_rel, gamma_rel***) were computed as the proportion of each band’s power to the total power.

The EDA raw signals were downsampled and smoothed to remove artifacts (Zhang et al., 2021). The signal was then decomposed into SCR and SCL, which represent the quickly changing phasic component and the slowly changing tonic component, respectively (Boucsein, 2012). Subsequently, the signals were segmented into 10-second epochs, and epochs with poor contact were excluded. For the remaining epochs, the average of SCL and the integrated SCR (iSCR) were computed. The means and standard deviations across epochs were calculated, resulting in the following metrics: SCL mean (***scl_mean***), SCL standard deviation (***scl_std***), iSCR mean (***iscr_mean***), and iSCR standard deviation (***iscr_std***).

For the PPG signal, the heart rate (***hr***) and pulse rate variability (***prv***) were estimated by identifying onsets of the waveform and computing the mean and standard deviation of pulse intervals (Kyriacou & Allen, 2021). To avoid interruptions caused by other periodic factors, epochs with heart rates outside the range of 50-100 bpm were excluded.

This study computed inter-subject correlations for EEG, SCR, and SCL signals. Pearson correlations were computed between each student’s signal and the average signal of other students (***eeg_isc, scr_isc, scl_isc***), as well as between the student and the teacher (***eeg_isc_t, scr_isc_t, scl_isc_t***).

The multimodal features are listed in Table 2. For all 777 10-minute sessions, there were 53, 67, and 25 samples with no valid EEG, EDA, and PPG epochs, respectively. Therefore, the missing features were interpolated using the mean value of other samples from the same student (Waljee et al., 2013).

**Table 2.**
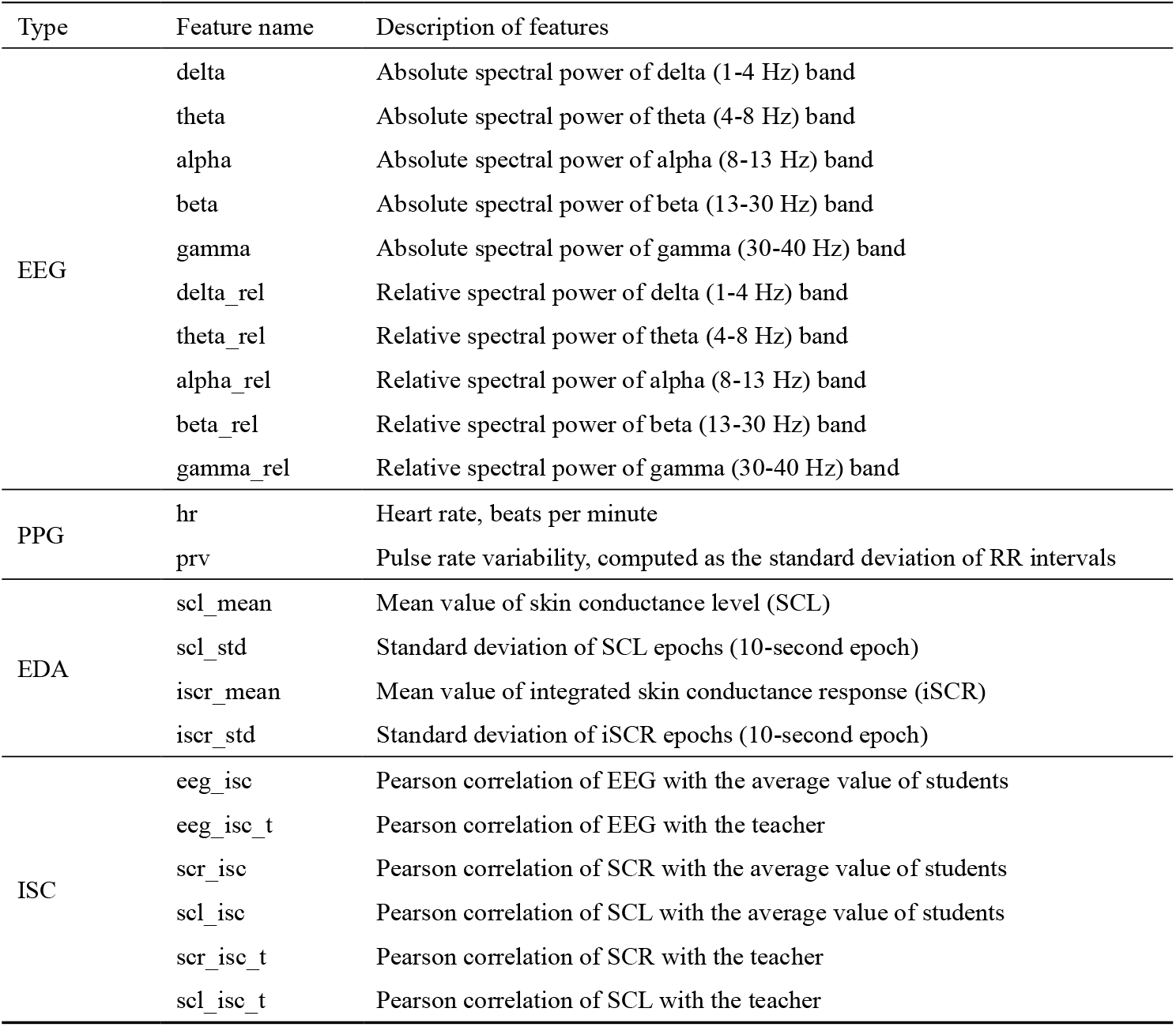
List of neurophysiological features.

## 4. Results

### 4.1 Descriptive statistics

First, the values of the cognitive and emotional variables for the high-performing and low-performing groups were examined. The average reported values were higher than 3 (neither agree nor disagree), indicating a positive learning status, as shown in Table 3. High-performing students gave slightly higher ratings than low-performing students across all variables. However, the group effect was not significant based on the estimation of equation (1).

**Table 3.** Descriptive statistics and group comparison.

| Variables | Descriptive statistics |  |  | Group differences |
| --- | --- | --- | --- | --- |
|  | All students | High-performing students | Low-performing students | High- and low-performing |
|  | (1) | (2) | (3) | (4) |
| Cognitive state | 3.701<br>[0.832] | 3.791<br>[0.805] | 3.600<br>[0.860] | 0.216<br>(0.291) |
| Cognitive regulation | 3.663<br>[0.904] | 3.740<br>[0.968] | 3.575<br>[0.817] | 0.209<br>(0.356) |
| Emotional state | 3.822<br>[0.880] | 3.915<br>[0.860] | 3.718<br>[0.892] | 0.280<br>(0.327) |
| Emotional regulation | 3.745<br>[0.921] | 3.818<br>[0.951] | 3.663<br>[0.879] | 0.238<br>(0.374) |
| N | 777 | 412 | 365 | 777 |

Note: Standard deviation in square brackets and standard error in parentheses. The group difference was estimated through equation (1).

The Pearson’s correlation coefficients were computed to analyze the relationship between cognition and emotion. Table 4 displays the pairwise correlations across all six combinations of variables, allowing for comparison across different subgroups. These correlation coefficients are all significant and show relatively large values. For the CS-ES and CR-ER variable pairs, the correlation coefficients are significantly higher in the ESM condition compared to the recall method. Moreover, within the ESM condition, the correlation coefficients are higher for high-performing students than for low-performing students (see Supplementary Note I for a detailed analysis).

**Table 4.**
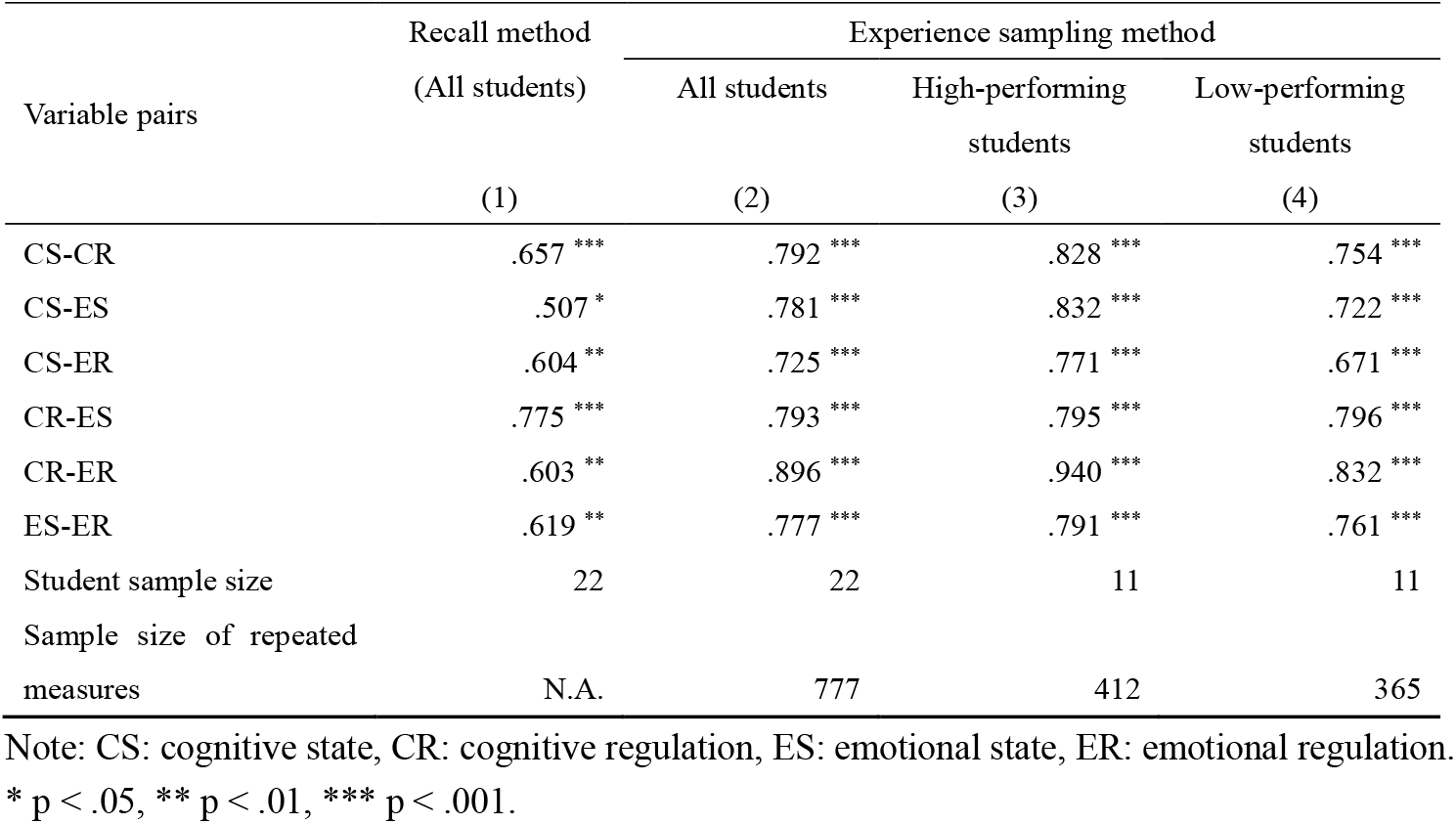
The Pearson’s correlation between the four variables.

| Variable pairs | Recall method | Experience sampling method |  |  |
| --- | --- | --- | --- | --- |
|  | (All students) | All students | High-performing students | Low-performing students |
|  | (1) | (2) | (3) | (4) |
| CS-CR | .657 *** | .792 *** | .828 *** | .754 *** |
| CS-ES | .507 * | .781 *** | .832 *** | .722 *** |
| CS-ER | .604 ** | .725 *** | .771 *** | .671 *** |
| CR-ES | .775 *** | .793 *** | .795 *** | .796 *** |
| CR-ER | .603 ** | .896 *** | .940 *** | .832 *** |
| ES-ER | .619 ** | .777 *** | .791 *** | .761 *** |
| Student sample size | 22 | 22 | 11 | 11 |
| Sample size of repeated measures | N.A. | 777 | 412 | 365 |

Note: CS: cognitive state, CR: cognitive regulation, ES: emotional state, ER: emotional regulation.
\* $p < .05$ , \*\* $p < .01$ , \*\*\* $p < .001$ .

### 4.2 Multilevel model analysis

The previous Pearson correlation analysis revealed high correlations between cognitive and emotional variables, as well as group differences. When analyzing through multilevel linear models, it was found that the differences in the cognitive-emotional relationship primarily stem from within-subject repeated-measures level, as shown in Table 5. The within-subject R-squared values for each dependent variable are significantly higher in high-performing students (ES: .610, ER: .791, CS: .663, CR: .793) than in low-performing students (ES: .181, ER: .132, CS: .171, CR: .140), highlighting a more intertwined relationship in high-performing students.

**Table 5.** The within, between, and overall R-squared values for the relationships between the variables.

| DV | All students |  |  | High-performing students |  |  | Low-performing students |  |  |
| --- | --- | --- | --- | --- | --- | --- | --- | --- | --- |
|  | Within | Between | Overall | Within | Between | Overall | Within | Between | Overall |
|  | (1) | (2) | (3) | (4) | (5) | (6) | (7) | (8) | (9) |
| CS | .461 | .822 | .675 | .663 | .838 | .746 | .171 | .806 | .600 |
| CR | .559 | .955 | .848 | .793 | .969 | .908 | .140 | .973 | .784 |
| ES | .458 | .865 | .704 | .610 | .834 | .728 | .181 | .903 | .690 |
| ER | .566 | .894 | .800 | .791 | .920 | .879 | .132 | .876 | .706 |

Note: The R-squared was computed based on the equation (2): $CS_{ij} = \beta_1 ES_{ij} + \beta_2 ER_{ij} + \beta_3 CR_{ij} + \mu_{0i} + \epsilon_{ij}$ . The four variables, where one serves as the dependent variable and the other three as independent variables, are alternated in this manner. DV: dependent variable, CS: cognitive state, CR: cognitive regulation, ES: emotional state, ER: emotional regulation.

### 4.3 Neurophysiological analysis

Figure 2 illustrates the neurophysiological representations of original values, fitted values, and residuals for the entire student sample. Significant associations were found between all the four original values and some neurophysiological features, such as ***hr, eeg_isc***, and ***eeg_isc_t***. These associations may result from a high proportion of shared components among the variables, as further confirmed by the analysis of fitted values. The fitted values (*y*_*hat*) of the four concepts (CS, CR ES, and ER) have common neurophysiological representations, including features such as ***delta_rel, hr, eeg_isc, eeg_isc_t***, and ***scr_isc***.

**Figure 2.**
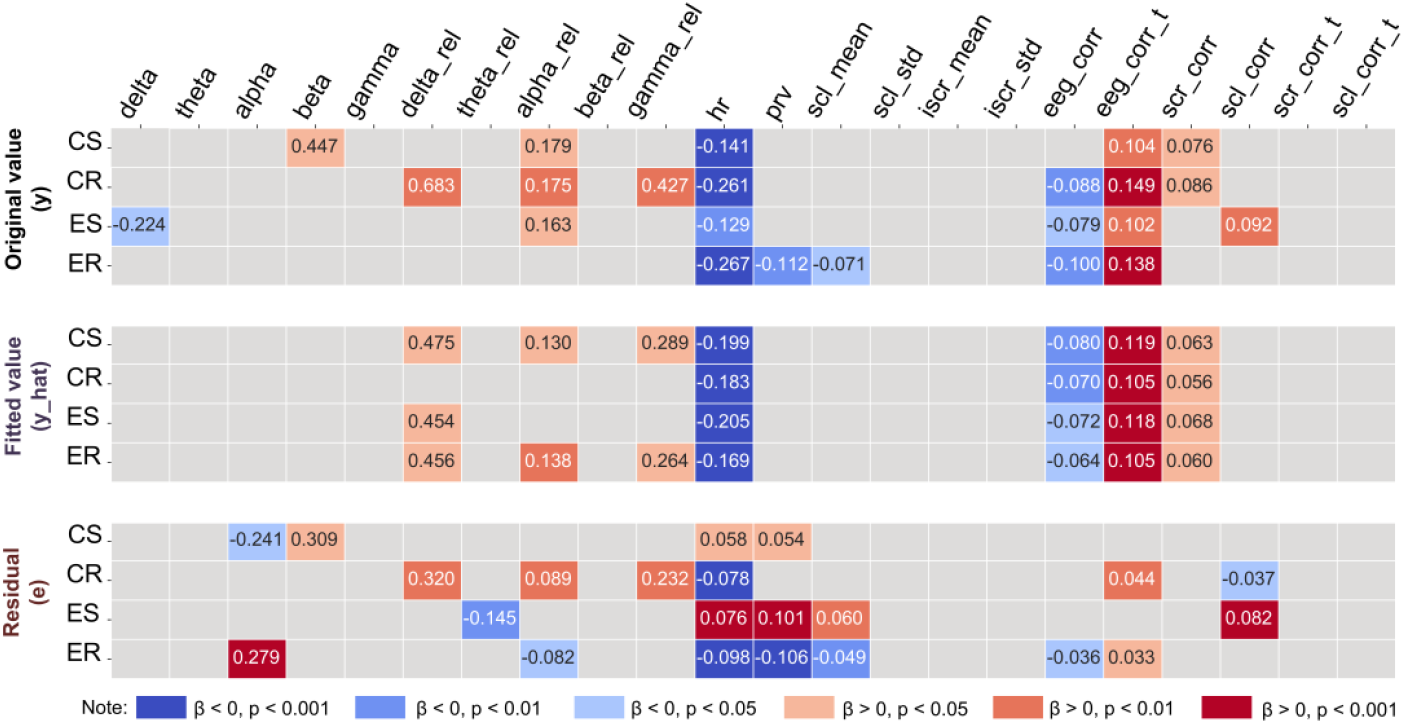
Neurophysiological representations of the raw values, fitted values, and residuals in the sample of all students. Coefficients were estimated using equation (4). Only significant coefficients are displayed in the figure.

**Figure 3.**
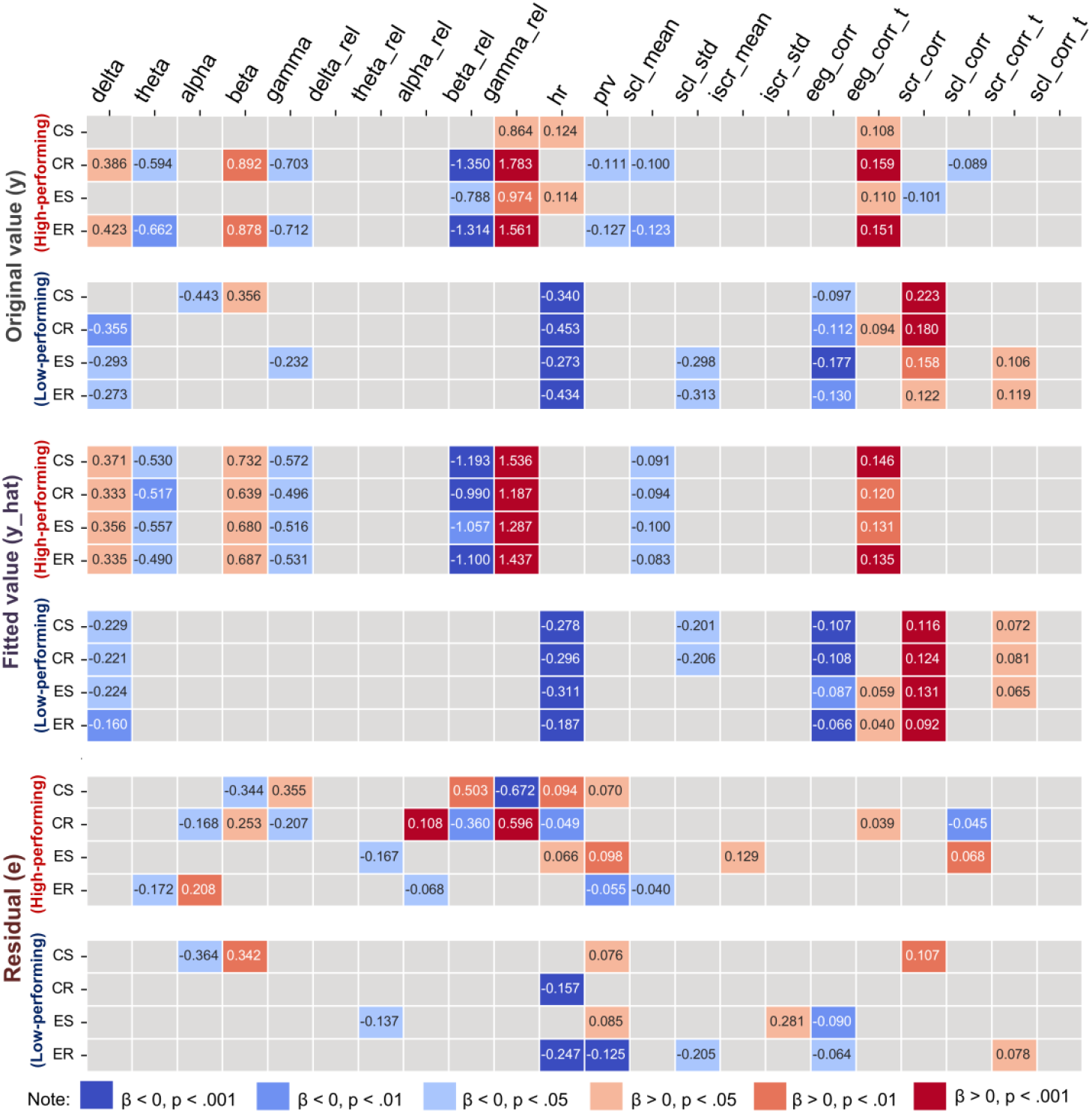
Neurophysiological representations of the raw values, fitted values, and residuals in the sample of high- and low-performing students. Coefficients were estimated using equation (4). Only significant coefficients are displayed in the figure.

The neurophysiological representations of the residuals (*e*) demonstrate significant differences, indicating unique neural mechanisms for cognition and emotion. For example, CS is linked to ***alpha, beta***, and ***prv***; CR is associated with ***scl_isc***; ES is associated with ***prv, scl_mean***, and ***scl_isc***; and ER is related to ***alpha, prv***, and ***scl_mean***. The neurophysiological analysis results of the fitted values and residuals align with our expectations, suggesting that the neurophysiological representations of cognition and emotion have both shared components and unique aspects.

When analyzing neurophysiological representations in high- and low-performing subgroups, group differences are particularly pronounced in the fitted values, which reflect the cognitive-emotional relationship. In high-performing students, the fitted values are positively significantly associated with ***delta, beta, gamma_rel***, and ***eeg_corr_t***; and negatively significantly associated with ***theta, gamma, beta_rel***, and ***scl_mean***. In low-performing students, significant features are quite different. The fitted values are positively significantly associated with ***scr_corr***; and negatively significantly associated with ***delta, hr***, and ***eeg_corr***. The analysis results suggest that the two groups may exhibit distinct cognitive and emotional patterns during learning.

## 5. Conclusion and Discussion

This study aims to investigate group differences in the cognitive-emotional relationship in a real-classroom learning environment. Utilizing experience sampling and multimodal data, the study explores the relationship between cognitive state, cognitive regulation, emotional state, and emotional regulation. Overall, results show that all four variables are strongly correlated, underscoring their interconnectedness in classroom learning. Regarding group differences, high-performing students showed stronger within-subject correlation among these factors than their lower-performing counterparts. This finding suggests that high achievers are more likely to maintain both heightened attention and positive emotions, with their regulation effectively supporting both. Furthermore, neurophysiological features showed significant associations with the fitted values of the four variables, with distinct representations for high- and low-performing students. The neurophysiological analysis supports the interplay between cognition and emotion, as well as the group differences in this relationship.

The high coefficients between cognition and emotion support the theoretical perspective that they are inseparable and mutually dependent. It is worth noting that the correlations identified in this study are comparable to, and in some cases numerically exceed those reported in previous literature, highlighting the unique characteristics of dynamic repeated-measures data. The r-values (on average .794, see Table 4 column (2)) exceed those observed in a dataset of high school students, where the cross-sectional r-values between emotional and cognitive engagement averaging .55 (Li & Lerner, 2013). These values are also higher than those between emotional and cognitive engagement reported in a MOOC learning dataset (on average .48) (Liu et al., 2022). Within the same student sample in this study, correlations are stronger in the ESM condition than in the full-scale survey (for detailed coefficient comparisons, see Supplementary Note I). While this high correlation might be under suspicion of being partly influenced by the experience sampling method, the observed differences between high- and low-performing students eliminate such concern. If the high correlation were solely due to the experience sampling instrument, such group differences would not be expected.

Although the cognitive-emotional relationship is generally strong, group differences are also evident, with high associations predominantly observed in high-performing students. A possible explanation is that the integration of cognition and emotion may indicate more effective learning. In educational theories, there has been growing interest in integrating emotions into the learning process, particularly regarding how emotions interact with cognition. For instance, Rogers, in his person-centered approach, highlighted the importance of “congruence” between cognition and emotion as a fundamental condition of learning and personal growth (Motschnig & Nykl, 2003). Vygotsky also argued that cognitive and emotional processes are inseparable. In his sociocultural theory, the integration of emotional and cognitive factors facilitates students’ progression toward their zone of proximal development (Mahn & John-Steiner, 2002). Based on these theories, it is expected that a high correlation between cognition and emotion is more likely to be observed in high-performing individuals.

The neurophysiological analysis provides a deeper explanation for the group differences in the cognitive-emotional relationship. Among high-performing students, the fitted values, which represent the shared components between cognition and emotion, are associated with multiple features. Some features have been shown to be related to effective learning. For example, the feature ***eeg_isc_t***, which is positively associated with the fitted values, represents student’s brain synchrony with the teacher. This feature indicates social interactions between teachers and students (Bevilacqua et al., 2019), and it has also been shown to causally predict learning outcomes (Xu et al., 2024). The teacher-student brain ISC was also found to exhibit temporal lag, suggesting that the knowledge transmission process occurs during these interactions (Zheng et al., 2018). Based on the discussion of the above literature, our findings suggest that effective cognitive-emotional interplay may be associated with high student-teacher interaction. Regarding single brain features, this study found that the fitted values were positively correlated with ***delta*** and ***beta*** band power and negatively correlated with ***theta*** and ***gamma*** band power. Both ***delta*** and ***beta*** features are associated with higher-order cognitive functions during learning processes (Harmony et al., 1996; Spitzer & Haegens, 2017). Additionally, the negative association with ***theta*** and the positive association with ***beta*** suggest that the fitted values are negatively associated with the theta-beta-ratio, as confirmed by statistical tests (see Supplementary Note V). The ratio has been regarded as a biomarker negatively correlated with focused attention (Putman et al., 2014; van Son et al., 2019). This finding suggests that effective cognitive and emotional collaboration may be linked to sustained concentration on the learning material.

Among low-performing students, neurophysiological representations differ substantially. For example, the feature ***hr*** is negatively associated with the fitted values. Heart rate (***hr***) is influenced by both the sympathetic and parasympathetic nervous systems, which means that both stimulus-induced arousal and regulatory processes influence its dynamics (Hughson et al., 1994; Solhjoo et al., 2019). The negative association with heart rate suggests that regulatory processes may play a dominant role in the relationship. The feature ***eeg_isc***, representing brain synchrony with the class average, also shows a negative correlation. Although the ISC with class average is considered a positive indicator of learning outcomes (Meshulam et al., 2021), the results of this study suggest that the brain activity of low-performing students was not aligned with the class average. In contrast, the feature ***scr_corr***, representing ISC with the class average based on SCR, shows a positive association with the fitted values. The SCR component reflects the rapid changes in skin response triggered by emotional or decision-making processes, and has been found to be negatively associated with academic performance (Zhang et al., 2018). Therefore, the ISC of the SCR signal suggests that these students may be experiencing similar emotional fluctuation.

In summary, this study adds empirical evidence to the longstanding debate on the relationship between emotion and cognition. Our findings support an overall strong correlation between cognition and emotion in real-world learning, with this relationship varying by academic performance level. This study also integrates multiple novel methods to enhance the validity of its conclusions. First, data collection in real-classroom settings enhances ecological validity compared to laboratory-based tasks. Second, experience sampling captures real-time feelings, reducing the effects of recall bias. Finally, multimodal data reveal the physiological basis for the connections between emotion and cognition, providing objective evidence for their relationship and group differences.

There are still several limitations that need improvement. First, there are many group difference factors that influence the relationship. This study only focuses on academic performance, while other factors, such as motivation, interest, grit, and self-regulation, also deserve consideration. Given the challenge of data collection, future research could include larger sample sizes and more diverse variables to better understand how learner characteristics influence the relationship between cognition and emotion. Second, due to the limited length of the experience sampling method questionnaires, both states and regulation could only be measured using very generalized items. Future studies could further investigate the specific relationships between different types of cognitive and emotional processes, identifying which ones are more beneficial for learning.

## Ethics Statement

The studies involving human participants were reviewed and approved by the Institutional Review Board (IRB) of the Department of Psychology, Tsinghua University.

## Acknowledgement

This study was supported by the National Natural Science Foundation of China (62177030) and the Beijing Educational Science Foundation of the Fourteenth 5-year Planning (BGEA23019). The authors would like to appreciate the assistance of Xinqiao Gao, Baosong Li, Xin Liu, Ziyang

Luo, Fei Qin, Yun Long, Haifeng Luo, Zhilin Qu, Guannan Yao, Zheng Dong, Ziyan Xu, Wanyan Sun, Xiaomeng Xu, Xuming Li, and Mingxuan Gao during the data collection.

