## Supplementary Materials for "Achievement-Based Differences in Cognitive-Emotional Interplay During Classroom Learning: A Multimodal Analysis with Experience Sampling Method"

**Supplementary Note I.** Statistical test of the Pearson's correlation.

For two Pearson's coefficient values, $r_1$ and $r_2$, the difference in their magnitudes can be tested by constructing a test statistic $Z$, as shown in equation (5):

$$Z = \frac{s_2 - s_1}{SE} \tag{5}$$

where $s_2 = \frac{1}{2}\ln\left(\frac{1+r_2}{1-r_2}\right)$, $s_1 = \frac{1}{2}\ln\left(\frac{1+r_1}{1-r_1}\right)$, $SE = \sqrt{\frac{1}{n_1-3} + \frac{1}{n_2-3}}$. When the value is sufficiently large, it indicates that $r_2$ is significantly higher than $r_1$. The statistical tests were conducted between the experience sampling method (ESM) and recall method surveys, as well as the ESM surveys between high- and low-performing students (see Supplementary Table S1).

**Supplementary Table S1.** Statistical tests of the Pearson coefficients for different columns in Table 4.

| | ESM vs. Recall method (columns (1) and (2)) | | | ESM high-performing vs. ESM low-performing (columns (3) and (4)) | | |
|---|---|---|---|---|---|---|
| | CS | CR | ES | CS | CR | ES |
| CR | 1.212 | | | 2.697 ** | | |
| ES | 2.052 * | 0.196 | | 3.909 *** | -0.040 | |
| ER | 0.922 | 3.156 ** | 1.320 | 2.914 ** | 7.520 *** | 1.045 |

Note: * $p < .05$, ** $p < .01$, *** $p < .001$. The column (x) in the table corresponds to the column number in Table 4. ESM: experience sampling method, CS: cognitive state, CR: cognitive regulation, ES: emotional state, ER: emotional regulation.

**Supplementary Note II.** Multilevel analysis on the relationship between variables.

In addition to the Pearson analysis, considering the hierarchical data structure, this study also used a multilevel model to analyze the pairwise correlations between variables, as shown in equation (6):

$$DV_{ij} = \beta_0 + \beta_1 IV_{ij} + \mu_{0i} + \epsilon_{ij} \quad (6)$$

where $DV_{ij}$ and $IV_{ij}$ are the dependent variable and the independent variable. For the four variables, a total of $A_4^2$ equations can be formed. Since swapping the independent and dependent variables does not affect the $R^2$ results, only the results of $C_4^2$ pairs were presented in Supplementary Table S2, including within-subject, between-subject, and overall R-squared.

**Supplementary Table S2.** Multilevel analysis on the relationship between variables.

| Items | All students | | | High-performing students | | | Low-performing students | | |
|---|---|---|---|---|---|---|---|---|---|
| | Within | Between | Overall | Within | Between | Overall | Within | Between | Overall |
| | (1) | (2) | (3) | (4) | (5) | (6) | (7) | (8) | (9) |
| Panel A: Structure of variance | | | | | | | | | |
| CS | 0.427 | 0.279 | 0.705 | 0.509 | 0.235 | 0.744 | 0.323 | 0.317 | 0.640 |
| CR | 0.638 | 0.204 | 0.842 | 0.603 | 0.132 | 0.734 | 0.652 | 0.268 | 0.920 |
| ES | 0.541 | 0.298 | 0.840 | 0.680 | 0.214 | 0.894 | 0.369 | 0.372 | 0.742 |
| ER | 0.707 | 0.202 | 0.909 | 0.783 | 0.113 | 0.896 | 0.606 | 0.280 | 0.886 |
| Panel B: R-squared between variables | | | | | | | | | |
| CS-CR | .366 | .812 | .627 | .617 | .775 | .684 | .075 | .864 | .569 |
| CS-ES | .330 | .822 | .610 | .498 | .896 | .693 | .123 | .774 | .522 |
| CS-ER | .340 | .668 | .526 | .569 | .645 | .594 | .061 | .675 | .450 |
| CR-ES | .328 | .822 | .629 | .510 | .756 | .631 | .074 | .916 | .633 |
| CR-ER | .488 | .918 | .802 | .752 | .942 | .884 | .073 | .906 | .693 |
| ES-ER | .369 | .780 | .604 | .565 | .695 | .625 | .076 | .842 | .578 |

Note: CS: cognitive state, CR: cognitive regulation, ES: emotional state, ER: emotional regulation. The R-squared was computed based on the equation (5).

For the results in equation (2) (see Table 5), Supplementary Table S3 provides the regression coefficients, which reflect how the variables influence each other in the regression model.

**Supplementary Table S3.** The coefficients of the equation (2)

| | All students | | | | High-performing students | | | | Low-performing students | | | |
|---|---|---|---|---|---|---|---|---|---|---|---|---|
| IV | CS | CR | ES | ER | CS | CR | ES | ER | CS | CR | ES | ER |
| CS | | .202 | .284 | .113 | | .217 | .344 | .064 | | .185 | .250 | .099 |
| CR | .395 | | .225 | .485 | .594 | | .093 | .674 | .249 | | .223 | .235 |
| ES | .300 | .120 | | .204 | .299 | .032 | | .184 | .306 | .207 | | .171 |
| ER | .214 | .520 | .397 | | .118 | .620 | .528 | | .188 | .406 | .288 | |
| cons | .010 | .010 | -.006 | -.021 | -.005 | .004 | .013 | -.003 | .024 | .012 | -.028 | -.054 |

Note: CS: cognitive state, CR: cognitive regulation, ES: emotional state, ER: emotional regulation. The regression was based on the equation (2): $CS_{ij} = \beta_1 ES_{ij} + \beta_2 ER_{ij} + \beta_3 CR_{ij} + \mu_{0i} + \epsilon_{ij}$.

**Supplementary Note III.** The within-subject scatter plot of the variables.

To provide a more intuitive illustration of the cognitive-emotional interplay, we created scatter plots for each pairwise correlation for every student, as shown in Supplementary Figure S1. For low-performing students (blue), the correlation coefficients vary, with some showing near-zero or even negative values. In contrast, most high-performing students (red) display positive correlations. Pearson correlation coefficients were compared between groups using t-tests, revealing that high-performing students had significantly higher correlations for some item pairs (ES-ER: t = 2.488, p = .026; ES-CS: t = 1.388, p = .200, ES-CR: t = 2.556, p = .024, ER-CS: t = 2.794, p = .014, ER-CR: t = 3.502, p = .004, CS-CR: t = 1.881, p = .083), supporting the Pearson analysis in Table 4. (Note: CS: cognitive state, CR: cognitive regulation, ES: emotional state, ER: emotional regulation)

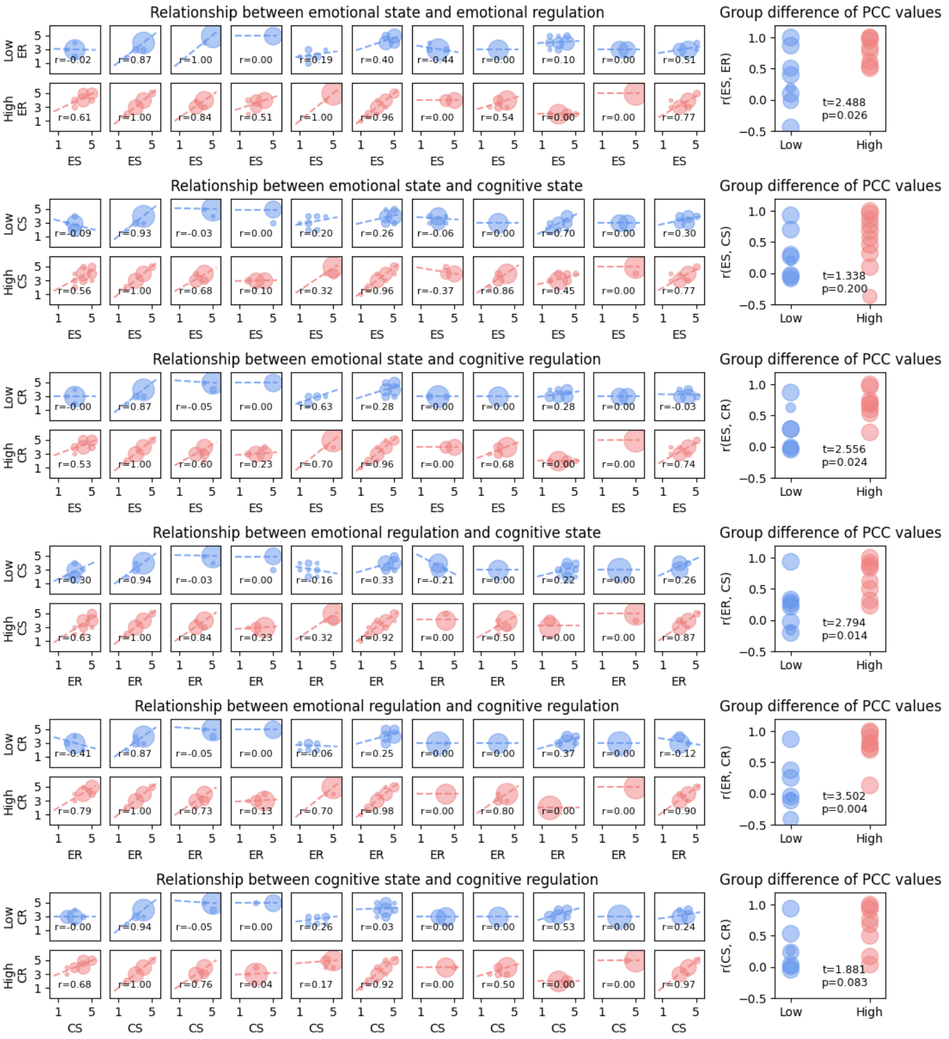


**Supplementary Figure S1.** Scatter plots of the cognitive and emotional variables for every student. For each pairwise correlation, we displayed the within-subject scatter plots for each individual in ascending order of student performance, arranged from top to bottom and left to right. We then examined the relationship between correlation coefficients and performance grouping.

Note: CS: cognitive state, CR: cognitive regulation, ES: emotional state, ER: emotional regulation, Low: low-performing students, High: high-performing students.

**Supplementary Note IV.** Further decomposition of the residual.

Based on the model in equation (2): $CS_{ij} = \beta_1 ES_{ij} + \beta_2 ER_{ij} + \beta_3 CR_{ij} + \mu_{0i} + \epsilon_{ij}$, the variance of the residuals ($\mu_{0i} + \epsilon_{ij}$) comprises two components: within-subject variance ($\epsilon_{ij}$, e_within) and between-subject variance ($\mu_{0i}$, e_between). These two components were also estimated using the model. Additionally, utilizing equation (4), neurophysiological representations for both components were derived, as shown in Supplementary Figure S2.

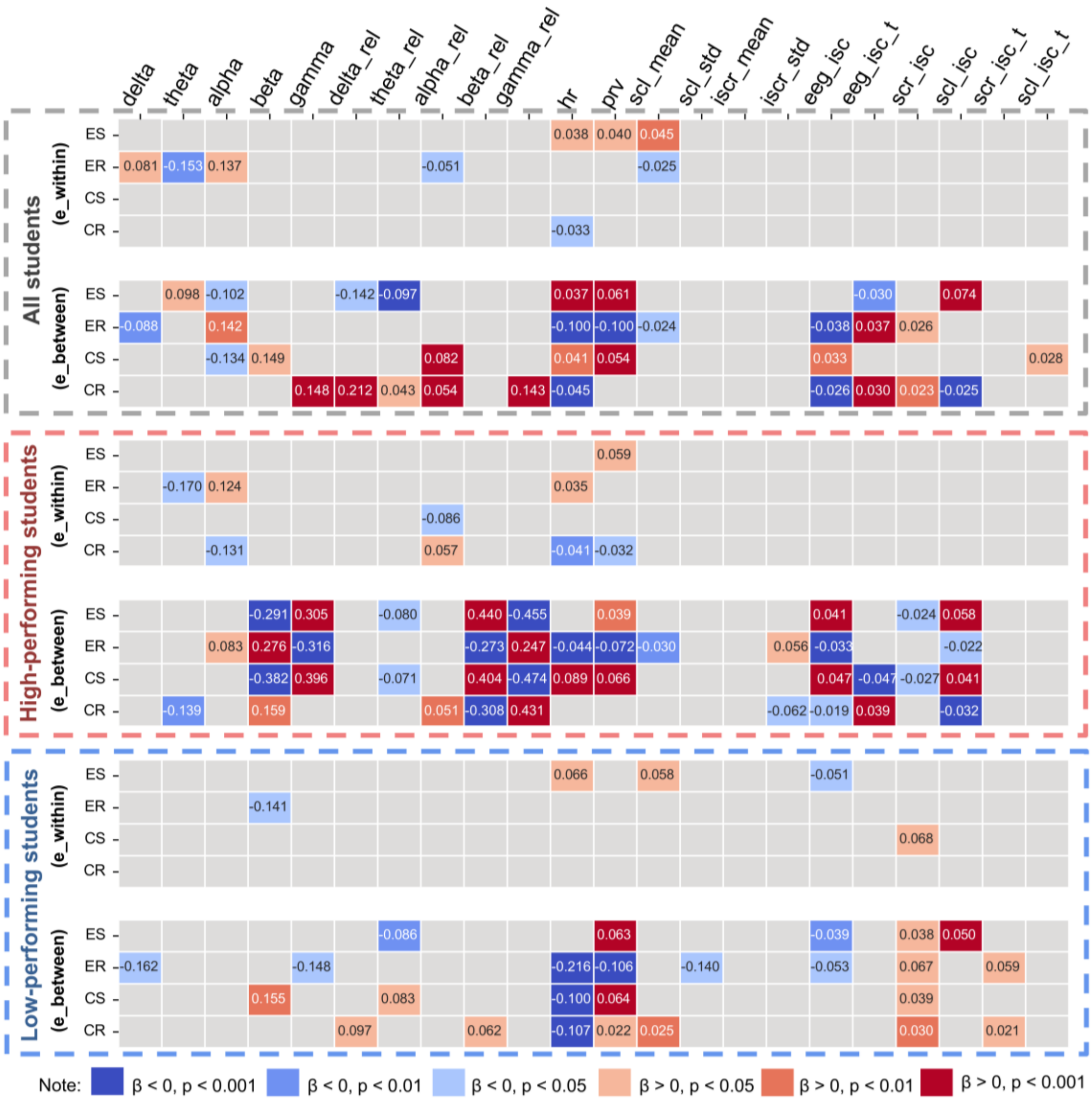


**Supplementary Figure S2.** Neurophysiological representations of within-subject and between-subject residuals. Coefficients were estimated through equation (4). Only significant coefficients are shown in the figure.

**Supplementary Note V.** Theta-beta-ratio as a potential feature.

When examining the neurophysiological representations of fitted values in high-performing students, this study identified a negatively associated feature, ***theta***, and a positively associated feature, ***beta***. This pattern suggests that the ***theta-beta-ratio***, a commonly used feature negatively correlated with focused attention, is likely negatively associated with the fitted values. Therefore, using equation (6), we explored the relationship between the ***theta-beta-ratio*** and the fitted values:

$$tbr = \beta_1 CS\_hat_{ij} + \mu_{0i} + \epsilon_{ij} \quad (6)$$

where $tbr = \frac{theta}{beta}$, representing the ratio between the band powers of theta and beta, and $CS\_hat_{ij}$ represents a specific fitted value (the fitted value of the cognitive state). The coefficients of equation (6) are presented in Supplementary Table S4.

**Supplementary Table S4.** The association between the theta-beta-ratio and the fitted values.

| Variable | $\beta_1$ | SE | p |
|---|---|---|---|
| CS_hat | -0.151 ** | 0.056 | .007 |
| CR_hat | -0.153 * | 0.067 | .022 |
| ES_hat | -0.174 ** | 0.060 | .004 |
| ER_hat | -0.146 * | 0.060 | .015 |

Note: CS: cognitive state, CR: cognitive regulation, ES: emotional state, ER: emotional regulation. The regression was based on the equation (6): $tbr = \beta_1 CS\_hat_{ij} + \mu_{0i} + \epsilon_{ij}$. * $p < .05$, ** $p < .01$

**Supplementary Note VI.** Gender differences on the cognitive-emotional relationship

Gender may also serve as an individual factor influencing learning. Therefore, in addition to analyzing the high- and low-performing groups, we also examined the cognitive-emotional relationship between genders using the same approach.

We first found that there were no significant differences in the data of cognitive and emotional variables between male and female students. Furthermore, the cognitive-emotional relationship was also examined for male or female participants, as shown in Supplementary Table S4. The results show that there are no significant differences in the $R^2$ values of each variable between female and male students, especially in terms of the within-group $R^2$. There is no noticeable difference between male and female students as pronounced as the difference between the high-performing and low-performing groups.

Neurophysiological analysis reveals potential gender differences. As shown in Supplementary Figure S3, the fitted values in girls are associated with ***theta_rel***, ***beta_rel***, ***gamma_rel***, and ***eeg_corr***, whereas in boys, these values are associated with ***hr*** and ***eeg_corr_t***. These analyses will help us understand the potential impact of gender on the relationship between cognition and emotion.

**Supplementary Table S5.** The within, between, and overall R-squared values for the relationships for girls or boys.

| | All students | | | Girls (9 students, 348 samples) | | | Boys (13 students, 429 samples) | | |
|---|---|---|---|---|---|---|---|---|---|
| DV | Within | Between | Overall | Within | Between | Overall | Within | Between | Overall |
| | (1) | (2) | (3) | (4) | (5) | (6) | (7) | (8) | (9) |
| CS | .461 | .822 | .675 | .323 | .786 | .626 | .547 | .890 | .752 |
| CR | .559 | .955 | .848 | .346 | .982 | .849 | .693 | .988 | .886 |
| ES | .458 | .865 | .704 | .304 | .855 | .665 | .585 | .886 | .751 |
| ER | .566 | .894 | .800 | .348 | .860 | .759 | .701 | .972 | .881 |

Note: The R-squared was computed based on the equation (2): $CS_{ij} = \beta_1 ES_{ij} + \beta_2 ER_{ij} + \beta_3 CR_{ij} + \mu_{0i} + \epsilon_{ij}$. The four variables, where one serves as the dependent variable and the other three as independent variables, are alternated in this manner. DV: dependent variable, CS: cognitive state, CR: cognitive regulation, ES: emotional state, ER: emotional regulation.

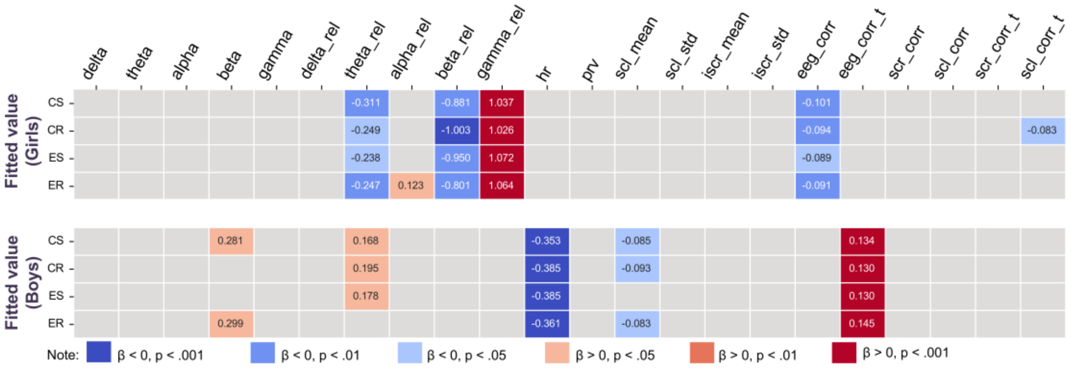


**Supplementary Figure S3.** Neurophysiological representations of the fitted values in the sample of girls or boys. Coefficients were estimated through equation (4). Only significant coefficients are shown in the figure.
